# Structural basis of anti-SARS-CoV-2 activity of HCQ: specific binding to N protein to disrupt its interaction with nucleic acids and LLPS

**DOI:** 10.1101/2021.03.16.435741

**Authors:** Mei Dang, Jianxing Song

## Abstract

Great efforts have led to successfully developing the spike-based vaccines but challenges still exist to completely terminate the SARS-CoV-2 pandemic. SARS-CoV-2 nucleocapsid (N) protein plays the essential roles in almost all key steps of the viral life cycle, thus representing a top drug target. Almost all key functions of N protein including liquid-liquid phase separation (LLPS) depend on its capacity in interacting with nucleic acids. Therefore, only the variants with their N proteins functional in binding nucleic acids might survive and spread in evolution and indeed, the residues critical for binding nucleic acids are highly conserved. Very recently, hydroxychloroquine (HCQ) was shown to prevent the transmission in a large-scale clinical study in Singapore but so far, no specific SARS-CoV-2 protein was experimentally identified to be targeted by HCQ. Here by NMR, we unambiguously decode that HCQ specifically binds NTD and CTD of SARS-CoV-2 N protein with Kd of 112.1 and 57.1 μM respectively to inhibit their interaction with nucleic acid, as well as to disrupt LLPS essential for the viral life cycle. Most importantly, HCQ-binding residues are identical in SARS-CoV-2 variants and therefore HCQ is likely effective to them all. The results not only provide a structural basis for the anti-SARS-CoV-2 activity of HCQ, but also renders HCQ to be the first known drug capable of targeting LLPS. Furthermore, the unique structure of the HCQ-CTD complex decodes a promising strategy for further design of better anti-SARS-CoV-2 drugs from HCQ. Therefore, HCQ is a promising candidate to help terminate the pandemic.

## Introduction

Severe Acute Respiratory Syndrome Coronavirus 2 (SARS-CoV-2) caused the ongoing catastrophic pandemic (1), which already led to >172 millions of infections and >3.7 millions of deaths. It belongs to a large family of positive-stranded RNA coronaviruses with ~30 kb genomic RNA (gRNA) packaged with nucleocapsid (N) protein into a membrane-enveloped virion. SARS-CoV-2 has four structural proteins: namely the spike (S) protein that recognizes the host-cell receptors angiotensin converting enzyme-2 (ACE2), membrane-associated envelope (E), membrane (M) proteins and N protein.

SARS-CoV-2 N protein is a 419-residue multifunctional protein (Fig. S1A), which is composed of the folded N-terminal domain (NTD) over residues 44-173 and C-terminal domain (CTD) over 248-365 (Fig. S1B), as well as three intrinsically-disordered regions (IDRs) respectively over 1-43, 174-247 and 366-419. Previous studies established that its NTD is an RNA-binding domain (RBD) functioning to bind various viral and host-cell nucleic acids including RNA and DNA (2), while CTD acts to dimerize/oligomerize to form high-order structures (3). Coronavirus N proteins appear to have two major categories of functions: while their primary role is to assemble gRNA and N protein to form the viral gRNA-Nprotein (vRNP) complex into the new virions at the final stage of the infection, they also act to suppress the immune system of the host cell and to hijack cellular machineries to achieve the replication of the virus, such as to interfere in the formation of stress granules (SGs) and to localize gRNA onto the replicase-transcriptase complexes (RTCs) (4-10).

Very recently, liquid-liquid phase separation (LLPS), the emerging principle for commonly organizing the membrane-less organelles (MLOs) or compartments critical for cellular physiology and pathology (11–15), has been identified as the key mechanism underlying the diverse functions of SARS-CoV-2 N protein (5–10). Nevertheless, all identified functions of N proteins so far appear to be dependent on its capacity in binding various nucleic acids with both specific and non-specific sequences. LLPS of N protein is mainly driven by its multivalent interactions with nucleic acids, as N protein with nucleic acids completely removed lacks the intrinsic capacity in LLPS (9,10). In particular, although the detailed mechanism still remains poorly understood, the final package of the RNA genome into new virions certainly requires the complex but precise interaction between gRNA and N protein, which should be extremely challenging for SARS-CoV-2 with such a large RNA genome (~30 kb). In this context, any small molecules capable of intervening in the interaction of N protein with nucleic acids are anticipated to significantly modulate most, if not all, key steps of the viral life cycle, some of which may thus manifest the anti-SARS-CoV-2 activity.

Hydroxychloroquine (HCQ), an antimalarial drug (Fig. 1C), has been extensively proposed for clinically combating the SARS-CoV-2 pandemic (16–18). Particularly, a large-scale clinical study in Singapore showed that oral HCQ can indeed prevent the spread of SARS-CoV-2 in the high transmission environments (18). However, the exact mechanisms for the anti-SARS-CoV-2 activity of HCQ remain highly elusive and so far, all the proposed action sites for HCQ are on the host cells, which include the interference in the endocytic pathway, blockade of sialic acid receptors, restriction of pH mediated S protein cleavage at the ACE2 binding site and prevention of cytokine storm (16,17). In particular, no SARS-CoV-2 protein has been experimentally identified to be targeted by HCQ.

**Figure 1.**
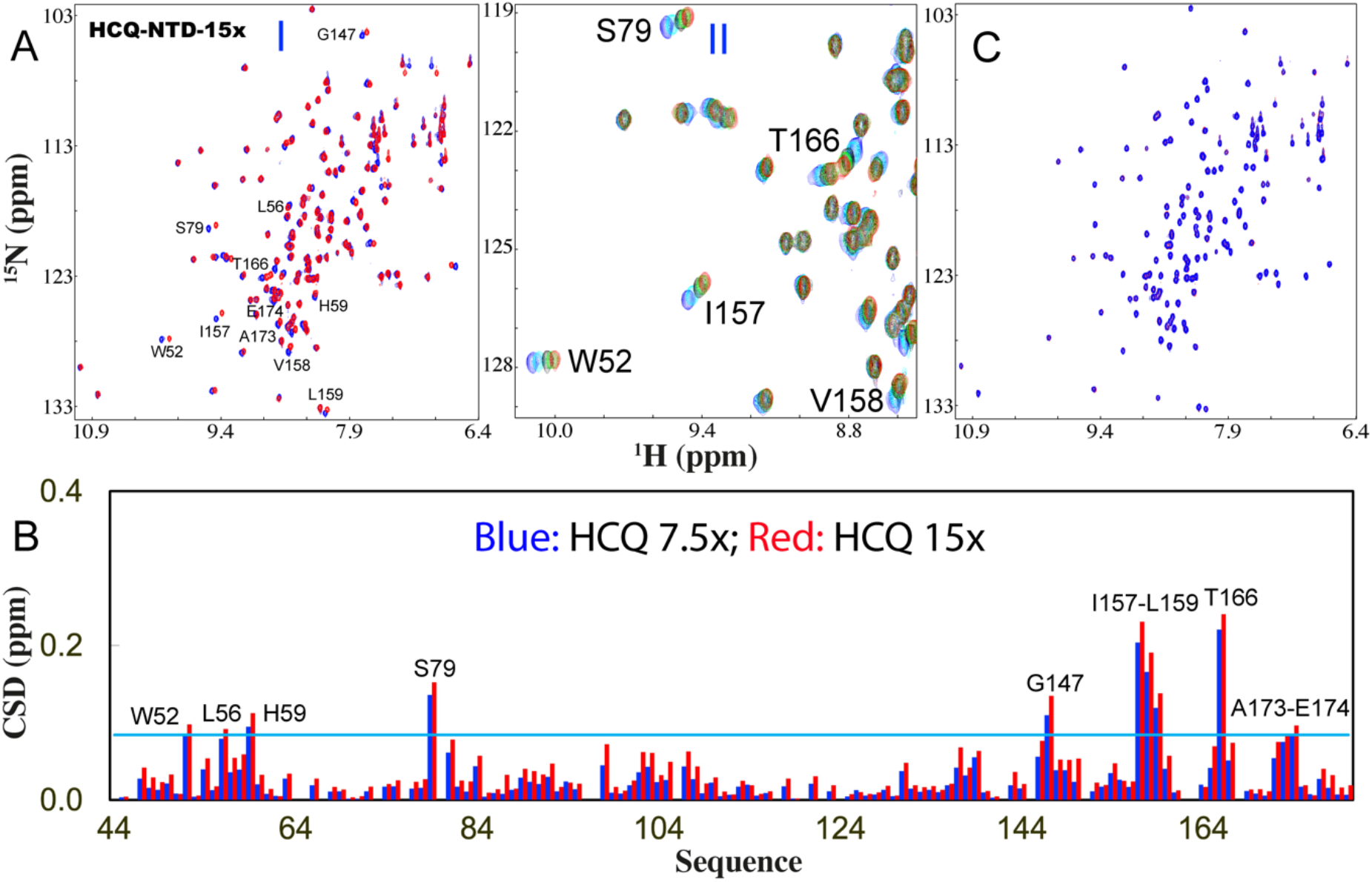
HCQ specifically binds NTD to inhibit its interaction with nucleic acid. (A) (I) Superimposition of HSQC spectra of NTD in the free state (blue) and in the presence of HCQ at 1:15 (NTD:HCQ) (red). The assignments of the significantly perturbed residues are labeled. (II) Zoom of HSQC spectra of NTD in the absence (blue) and in presence of HCQ at 1:1.88 (cyan); 1:3.75 (black); 1:7.5 (green); 1:15 (red). (B) Residue-specific chemical shift difference (CSD) of NTD in the presence of HCQ at 1:7.5 (blue) and 1:15 (red) (NTD:HCQ). The significantly perturbed residues are labeled, which are defined as those with the CSD values at 1:15 > 0.084 (average value + one standard deviation) (cyan line). (C) Superimposition of HSQC spectra of NTD only in the presence of HCQ at 1:15 (red) and with additional addition of S2m at 1;2.5 (NTD:S2m). (blue)

Very unexpectedly, here our residue-specific characterization by NMR spectroscopy decrypted that: 1) in addition to NTD, CTD can also bind nucleic acids with the affinity even higher than that of NTD. 2) HCQ specifically binds both NTD and CTD with Kd of 112.1 and 57.1 μM respectively to inhibit their interactions with nucleic acids. 3) HCQ has no capacity in inducing LLPS of N protein but is able to dissolve LLPS of N protein induced by nucleic acids, which is underlying the key steps of the viral life cycle. Therefore, the results not only provide a structural basis for the anti-SARS-CoV-2 activity of HCQ, but to the best of our knowledge, render HCQ to be the first known drug capable of targeting LLPS. Furthermore, the unique structure of the HCQ-CTD complex offers a promising strategy for further design of better anti-SARS-CoV-2 drugs from HCQ.

## Results

### HCQ specifically bind NTD to inhibit its interaction with nucleic acid

Very recently, NTD of SARS-CoV-2 N protein was determined by NMR and HADDOCK to utilize a conserved surface to bind various nucleic acids of random sequences or derived from the viral gRNA with dissociation constant (Kd) of ~ μM (2). Here, we first aimed to establish a NTD-nucleic acid binding system by titrating the ^15^N-labeled NTD sample with a 32-mer stem-loop II motif (S2m) single-stranded DNA (ssDNA) derived from the SARS-CoV-2 viral gRNA (Fig. S1C), which is a highly conserved sequence among coronaviruses and has been extensively utilized to characterize the nucleic-acid-binding of not only NTD, but also CTD of SARS-CoV-1 (4).

NMR spectroscopy is a very powerful tool for the residue-specific characterization of the weak binding associated with instable and aggregation-prone protein samples, which are thus not amenable to investigations by other biophysical methods such as ITC (19–21). Indeed, as monitored by NMR, S2m could bind NTD characteristic of broadening of HSQC peaks (Fig. S2). At 1:2.5 (NTD:S2m), a large portion of HSQC peaks became disappeared and further addition of S2m led to no significant change of HSQC spectra. This observation implies that the binding of NTD with S2m is over the intermediate NMR time scale (19–21), consistent with the recent report (2).

Subsequently, we evaluated the binding of HCQ to NTD by stepwise addition of HCQ as monitored by NMR. Strikingly, only a small set of HSQC peaks were shifted and the shifting process was largely saturated at 1:15 (NTD:HCQ) (Fig. 1A), unambiguously revealing that HCQ does specifically bind NTD. All NMR HSQC spectra in the presence of HCQ at different ratios were assigned and only eleven residues including Trp52, Leu56, His59, Ser79, Gly147, Ile157, Val158, Leu159, Thr166, Ala173 and Glu174 were significantly perturbed upon adding HCQ (Fig. 1B). Although these residues are distributed over the whole NTD sequence, they become clustered together in the NTD structure (Fig. 2A). We then conducted fitting of the NMR shift tracings by the well-established procedure (2,10,19,22) of 11 residues to obtain their residue-specific Kd values (Table S1 and Fig. 2B) with the average Kd of 112.1 ± 32.2 μM. Furthermore, with NMR-derived constraints, the structure of the HCQ-NTD complex was constructed (Fig. 2C) by the well-established HADDOCK program (20–23). In the complex structure, the binding pocket of HCQ is located within the conserved surface for NTD to bind various nucleic acids (2).

**Fig. 2.**
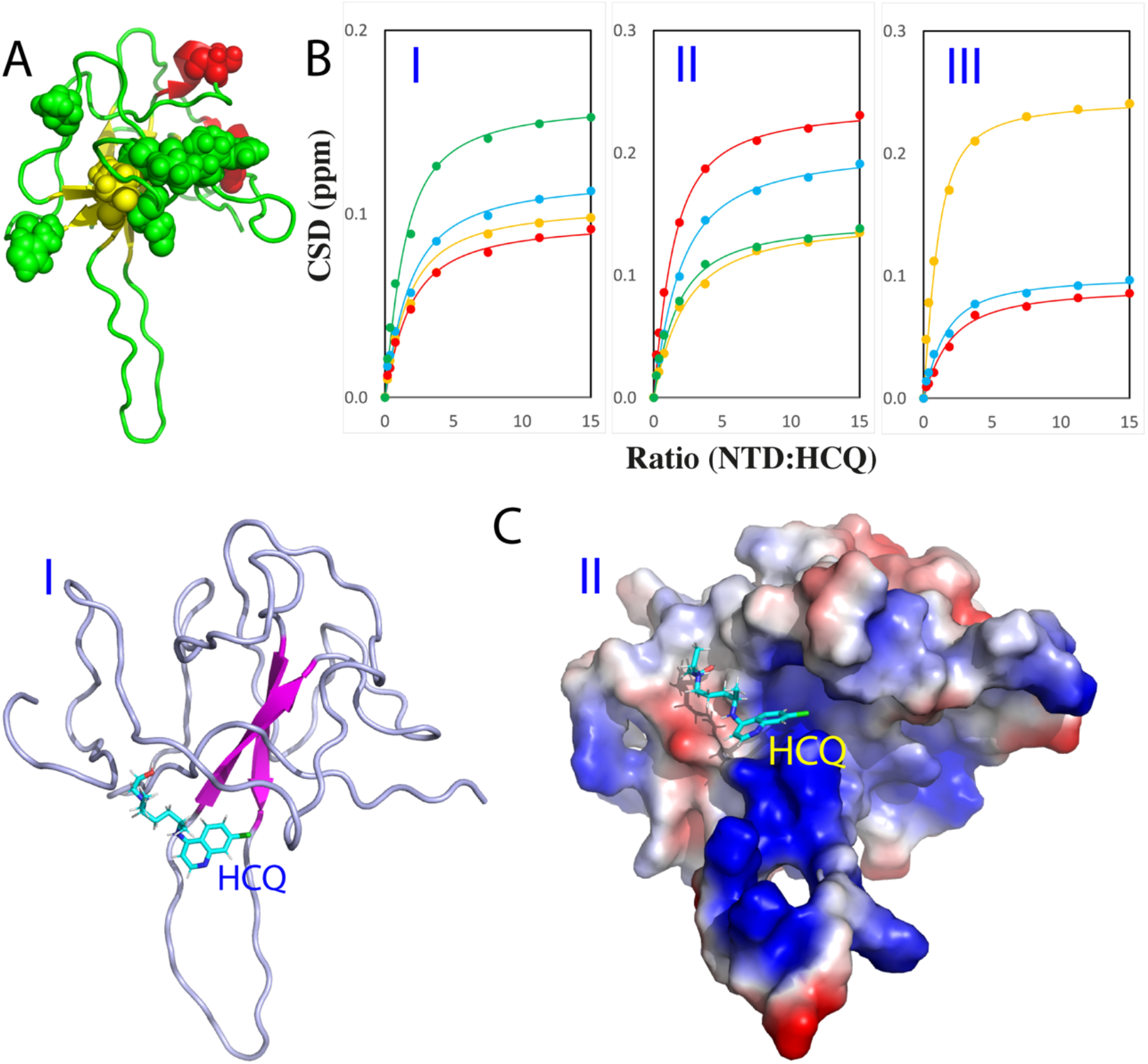
NMR characterization of the binding of HCQ to NTD. (A) 11 significantly perturbed residues of NTD upon binding to HCQ. (B) Fitting of 11 NTD residuespecific dissociation constant (Kd): experimental (dots) and fitted (lines) values for the CSDs induced by addition of HCQ at different ratios. (I) Trp52 (brown), Leu56 (red), His59 (cyan), Ser79 (green). (II) Gly147 (brown), Ile157 (red), Val158 (cyan), Leu159 (green). (III) Thr166 (brown), Ala173 (red), Glu174 (cyan). (C) Structure of the HCQ-NTD complex with HCQ in sticks and NTD in ribbon (I) and in electrostatic potential surface (II).

We then asked a question whether the HCQ binding can interfere in the binding of NTD with S2m. To address this question, we added S2m to the NTD sample with the pre-existence of HCQ at 1:15 and found no significant change of HSQC spectra even up to 1:2.5 (NTD:S2m) (Fig 1C). This result suggests that the HCQ-bound NTD was inhibited for further binding to S2m. To confirm this, we also prepared the NTD sample pre-saturated with S2m at 1:2.5 and then stepwise added HCQ into this sample (Fig. S3). Interestingly, even upon adding HCQ at 1:3.75 (NTD:HCQ), many disappeared HSQC peaks due to being bound with S2m became restored and at 1:15, all HSQC peaks could become detected which are very similar to those of the NTD sample only in the presence of HCQ at 1:15 (NTD:HCQ). This result clearly indicates that HCQ is even able to displace S2m from being bound with NTD.

### HCQ specifically bind CTD to inhibit its interaction with nucleic acid

In parallel, we titrated the ^15^N-labeled CTD with S2m and unexpectedly S2m was also able to bind CTD characterized by broadening of HSQC peaks (Fig. S4). Most HSQC peaks became disappeared even at 1:1 (CTD:S2m) and further addition of S2m induced no significant change. This is similar to what was previously observed on SARS-CoV-1 N protein: its CTD binds S2m with the affinity higher than that of NTD (4).

Subsequently, we stepwise added HCQ into the CTD sample and again only a small set of HSQC peaks became shifted. At 1:7.5 (CTD:S2m) the shifting process was largely saturated (Fig. 3A). Detailed analysis of the spectra at different HCQ concentrations revealed that only seven CTD residues including Gln281, Thr282, Thr325, Thr329, Trp330, Ala336 and Ile337 were significantly perturbed upon adding HCQ (Fig. 3B), which are located over the whole sequence but clustered together to form two pockets in the dimeric CTD structure (Fig. 4A). The residue-specific Kd values of seven residues were successfully obtained (Table S1 and Fig. 4B) with the average Kd of 52.1 ± 17.6 μM. The structure of the HCQ-CTD complex was also constructed from NMR-derived constraints in which the dimeric CTD is bound with two HCQ molecules (Fig. 4C). Interestingly, the binding of HCQ appears to be mainly driven by the insertion of the aromatic rings of HCQ into the pockets of the cleft on the same side of the dimeric CTD structure.

**Figure 3.**
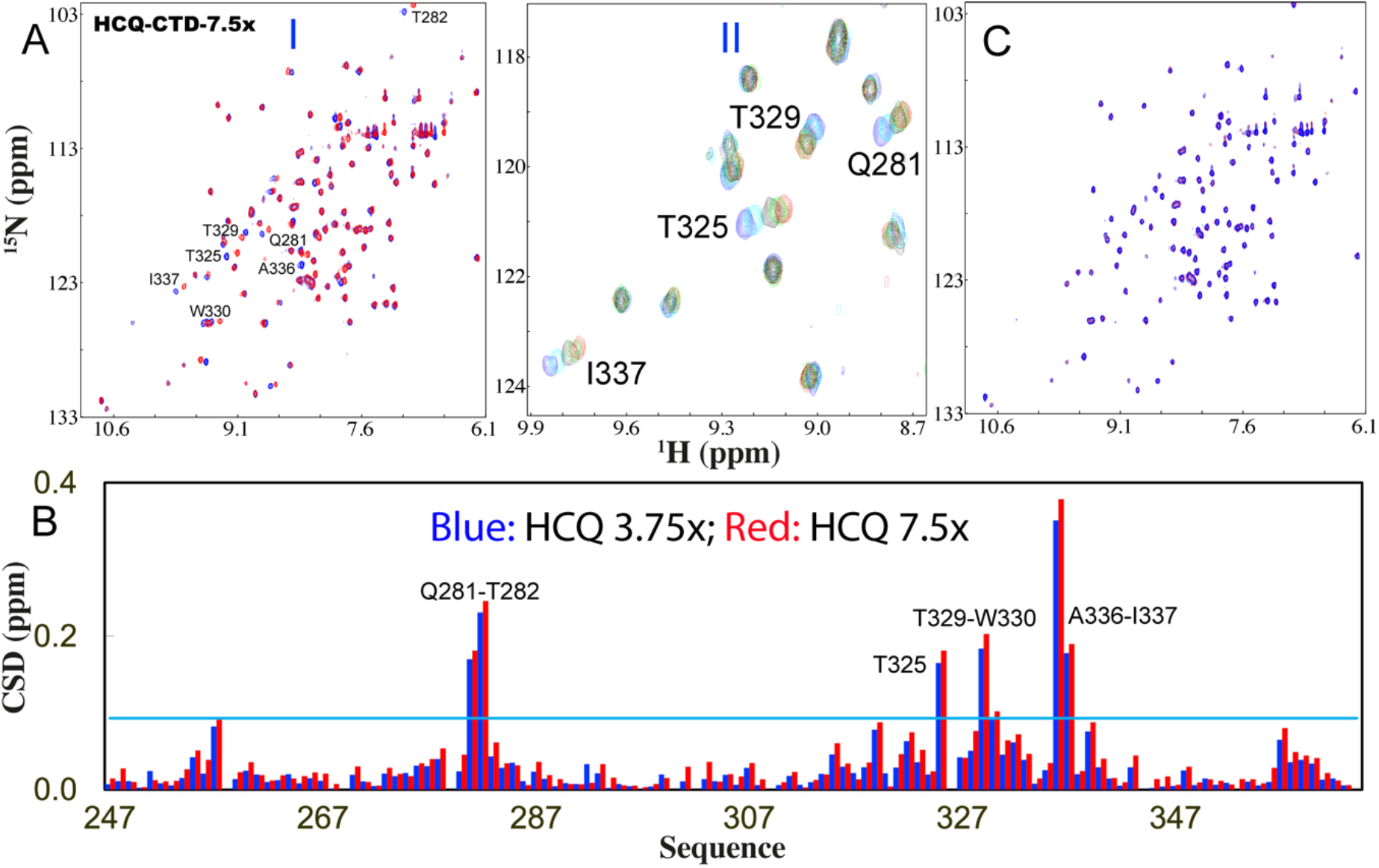
HCQ specifically binds CTD to inhibit its interaction with nucleic acid. (A) (I) Superimposition of HSQC spectra of CTD in the free state (blue) and in the presence of HCQ at 1:7.5 (CTD:HCQ) (red). (II) Zoom of HSQC spectra of CTD in the absence (blue) and in presence of HCQ at 1:0.925 (cyan); 1:1.86 (black); 1:3.75 (green); 1:7.5 (red). The assignments of the significantly perturbed residues are labeled. (B) Residue-specific chemical shift difference (CSD) of CTD in the presence of HCQ at 1:3.75 (blue) and 1:7.5 (red) (CTD:HCQ). The significantly perturbed residues are labeled, which are defined as those with the CSD values at 1:7.5 > 0.093 (average value + one standard deviation) (cyan line). (C) Superimposition of HSQC spectra of CTD only in the presence of HCQ at 1:7.5 (red) and with additional addition of S2m at 1;1 (CTD:S2m) (blue).

**Fig. 4.**
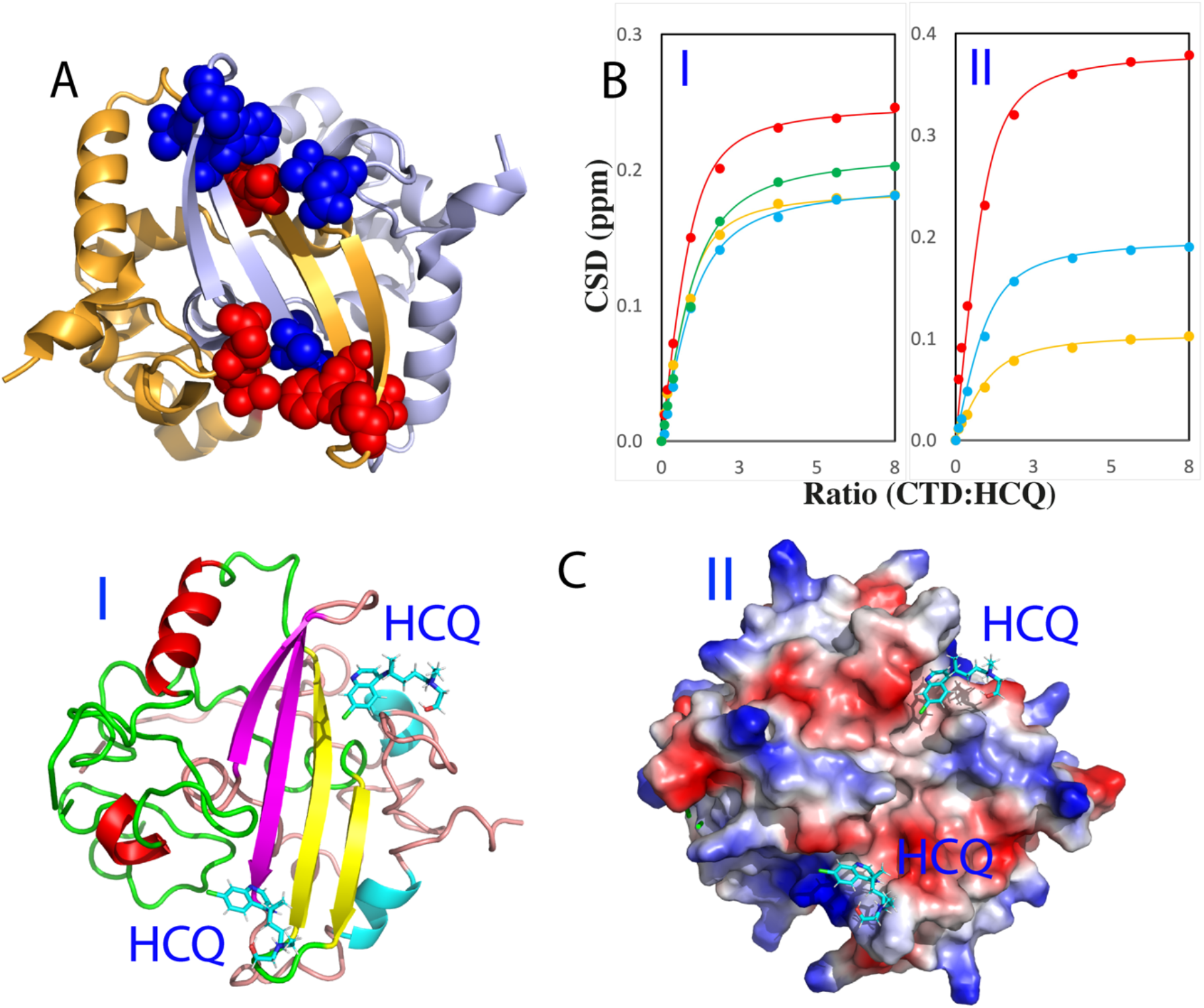
NMR characterization of the binding of HCQ to CTD. (A) 7 significantly perturbed residues of CTD upon binding to HCQ. (B) Fitting of 7 CTD residue-specific dissociation constant (Kd): experimental (dots) and fitted (lines) values for the CSDs induced by addition of HCQ at different ratios. (I) Gln281 (brown), Thr282 (red), Thr325 (cyan), Thr329 (green). (II) Trp330 (brown), Ala336 (red), Ile337 (cyan). (C) Structure of the HCQ-CTD complex with HCQ in sticks and CTD in ribbon (I) and in electrostatic potential surface (II).

We also assessed the interplay of HCQ and S2m in binding CTD. Briefly, we added S2m to the CTD sample in the pre-existence of HCQ at 1:7.5 but found no significant change of HSQC spectra even up to 1:1 (CTD:S2m) (Fig. 3C). This observation suggests that the HCQ-bound CTD is also blocked for binding with S2m. We also prepared the CTD sample presaturated with S2m at 1:1 into which HCQ was stepwise added (Fig. S5). Interestingly, upon adding HCQ at 1:1.8 (NTD:HCQ), many disappeared HSQC peaks became restored and at 1:7.5, all HSQC peaks could become detected which are highly similar to those of the CTD sample only in the presence of HCQ at 1:7.5 (CTD:HCQ). Although so far, the efforts failed to determine the structures of CTD of either SARS-CoV-1 or SARS-CoV-2 in complex with nucleic acids, here the NMR results unambiguously indicates that HCQ can also displace S2m from being bound with CTD.

### HCQ dissolves LLPS of N protein induced by nucleic acid

Very recently, the SARS-CoV-2 N protein was shown to function through LLPS, which was induced by dynamic and multivalent interactions with various nucleic acids including viral and host-cell RNA and DNA (5–10). Here we thus asked a question whether HCQ has any effect on LLPS of SARS-CoV-2 N protein. To address this, we first titrated S2m into the fulllength N protein and imaged LLPS by DIC microscopy as we previously conducted on SARS-CoV-2 N protein (10) and FUS (14). Consistent with the previous reports including by us with specific and non-specific nucleic acid fragments of different lengths (5–10), S2m also imposed the biphasic effect on LLPS of N protein: induction at low ratios and dissolution at high ratios. Briefly, while the N protein showed no LLPS in the free state, LLPS was induced upon addition of S2m as evidenced by turbidity and DIC imaging. At 1:0.75 (Nprotein:S2m), the turbidity reached the highest value of 1.92 (I of Fig. 5A) and many liquid droplets were formed (Fig. 5B). Further addition of S2m dissolved LLPS and at 1:1.5 all liquid droplets were dissolved.

**Figure 5.**
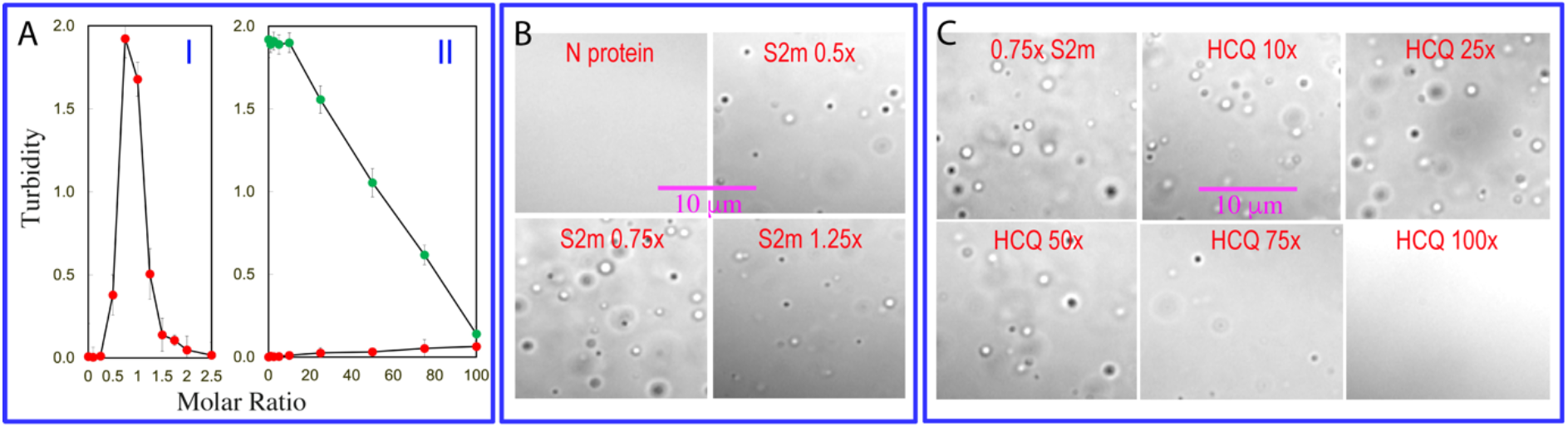
HCQ disrupts LLPS of SARS-CoV-2 N protein. (A) Turbidity curves of N protein in the presence of S2m at different ratios (I). (II) Turbidity curves of N protein in the absence (red) or in the presence of S2m at 1:0.75 (green) with additional addition of HCQ at different ratios. (B) DIC images of N protein in the presence of S2m at different ratios. (C) DIC images of N protein in the presence of S2m at 1:0.75 with additional addition of HCQ at different ratios.

We then stepwise added HCQ into the N protein sample under the same conditions but no LLPS occurred even with the ratio up to 1:100 (II of Fig. 5A), indicating that HCQ is incapable of inducing LLPS. We then prepared a phase separated N protein sample with the pre-presence of S2m at 1:0.75 and subsequently added HCQ into this sample in a stepwise manner, as monitored by turbidity (II of Fig. 5A) and DIC imaging (Fig. 5C). At the ratios <1:20, HCQ has no significant effect on LLPS. However, at ratios >1:20, HCQ monotonically dissolved liquid droplets as evidenced by gradual reduction of turbidity and disappearance of liquid droplets. Strikingly, at 1:100 (Nprotein:HCQ), liquid droplets were completely dissolved. Previously it was established that the dynamic and multivalent binding of nucleic acids to SARS-CoV-2 N protein plays a key role in driving its LLPS (5–10). Therefore, HCQ acts to dissolve LLPS by displacing the nucleic acid from being bound with both NTD and CTD as we revealed above.

## Discussion

So far, great efforts have been dedicated to successfully developing the spike-based vaccines to combat the pandemic. Nevertheless, many challenges still remain to completely terminate the pandemic, which include the rapidly-emerging antibody-resistance variants (24), the adverse effects of the spike protein (25) and even its antibody (26). Most seriously, SARS-CoV-2 spike protein has been identified to provoke antibody-dependent enhancement (ADE) of infection (27,28) while its RNA fragments were shown to get integrated into human genome (29). Therefore, any small molecule drugs that directly target SARS-CoV-2 proteins to disrupt its life cycle are extremely valueless and urgently demanded to finally terminate the pandemic.

Out of SARS-CoV-2 proteins, N protein is the only one which plays the essential roles in almost all key steps of the viral life cycle, thus representing a top drug target. Mechanistically, almost all functions of N protein including LLPS appear to depend on its capacity in interacting with a variety of nucleic acids of diverse sequences. As a consequence, in evolution only the SARS-CoV-2 variants with their N protein functional in binding nucleic acids can survive and spread. Indeed, as shown in Fig. S6, the sequences of NTD and CTD critical for binding nucleic acids are not only highly conserved in the variants of SARS-CoV-2, but also in SARS-CoV-1. In this context, any small molecules capable of blocking in the interaction of N protein with nucleic acids would disrupt the viral life cycle.

Here, for the first time, HCQ, a safe drug recommended by WHO to treat other diseases (16–18), has been decrypted to specifically bind NTD and CTD of SARS-CoV-2 N protein to inhibit its interactions with nucleic acids as well as to dissolve LLPS. This finding not only provided an acting mechanism for the anti-SARS-CoV-2 activity of HCQ, but also validated that SARS-CoV-2 N protein and its LLPS are indeed druggable by small molecules, thus opening up a promising avenue for further development of anti-SARS-CoV-2 drugs by targeting N protein. For example, previously it was shown that HCQ could inhibit the maturation of SARS-CoV-2 virions, but this was proposed to result from the HCQ-induced changes of host-cell structures/conditions such as pH (16,17). The current results suggest that the ability of HCQ to specifically disrupt LLPS of N protein induced by nucleic acids may at least partly contribute to the inhibition of the maturation of SARS-CoV-2 virions. To the best of our knowledge, HCQ is the first drug which has been revealed to target LLPS, thus bearing the unprecedented implications for further design of drugs in general by modulating LLPS, whose roles in underlying various human diseases are starting to be recognized (5–15).

At the fundamental level of molecular interactions, the core mechanisms for LLPS induced by nucleic acids appear to be relatively conserved (10,14,15). In particular, the significantly perturbed residues of NTD and CTD upon binding HCQ are completely identical in all variants of SARS-CoV-2, as well as even highly conserved in SARS-CoV-1 (Fig. S6). As a consequence, HCQ which acts to block the nucleic-acid-binding and LLPS of N proteins is anticipated to be effective to most, if not all, variants of SARS-CoV-2. On the other hand, emerging evidence suggests that the nucleic-acid-induced LLPS is not only essential for the life cycles of coronaviruses, but also might be generally critical for other virus-host interactions. This might thus explain the puzzling observations that HCQ was not only effective in treatment of SARS-CoV-2 and SARS-CoV-1, but also to Dengue/Zika infections (16,17). Moreover, in the future it would be also of significant interest to explore whether the mechanisms of HCQ in treating other diseases are related to its capacity in intervening in LLPS.

The relatively low binding affinities of HCQ to NTD and CTD of SARS-CoV-2 N protein might rationalize the reports that HCQ is effective in preventing the infection as well as in treating the infection at the early stage (16–18). Nevertheless, the unique HCQ-CTD structure (Fig. 4) in which two HCQ molecules use the aromatic rings to insert into two binding pockets of a cleft on the same side of the dimeric CTD might offer a promising strategy for further design of anti-SARS-CoV-2 drugs with better affinity and specificity. Briefly, by introducing the proper groups to link two HCQ aromatic rings, the bivalent or even multivalent binders might be obtained, whose Kd value is the time of the Kd values of individual groups (30). In particular, those bivalent/multivalent binders are also anticipated to have the high specificity because the probability for human proteins to have such unique two HCQ binding pockets of the dimeric CTD should be extremely low. In fact, we have already constructed such a molecule in silico designated as DiHCQ by linking two HCQ molecules (Fig. S7A) and conducted the docking to the dimeric CTD. Indeed, two aromatic rings of DiHCQ do bind the pockets in the manner similar to those of two individual HCQ molecules (Fig. S7B).

Due to the extreme urgency to combat the pandemic, here we propose to integrate the combinatorial chemistry as well as biophysical methods including experimental such as NMR (20-22,30,32) and computational such as MD simulations (31,32) to design bivalent or even multivalent small molecules starting from HCQ to obtain more efficient anti-SARS-CoV-2 drugs, which can bind the dimeric CTD of SARS-CoV-2 N protein not only by its two aromatic rings, but also by the linker groups.

## Materials and methods

### Preparation of recombinant SARS-CoV-2 nucleocapsid as well as its NTD and CTD

The gene encoding 419-residue SARS-CoV-2 N protein was purchased from a local company (Bio Basic Asia Pacific Pte Ltd), which was cloned into an expression vector pET-28a with a TEV protease cleavage site between N protein and N-terminal 6xHis-SUMO tag used to enhance the solubility. The DNA fragments encoding its NTD (44–180) and CTD (247-364) were subsequently generated by PCR rection and cloned into the same vector (10).

The recombinant N protein and its NTD/CTD were expression in *E. coli* cells BL21 with IPTG induction at 18 °C. Both proteins were found to be soluble in the supernatant. For NMR studies, the bacteria were grown in M9 medium with addition of (^15^NH_4_)_2_SO_4_ for ^15^N-labeling. The recombinant proteins were first purified by Ni^2+^-affinity column (Novagen) under native conditions and subsequently in-gel cleavage by TEV protease was conducted. The eluted fractions containing the recombinant proteins were further purified by FPLC chromatography system with a Superdex-200 column for the full-length and a Superdex-75 column for NTD and CTD (10). The purity of the recombinant proteins was checked by SDS-PAGE gels and NMR assignment for both NTD and CTD. Hydroxychloroquine (HCQ) sulfate was purchased from Merck (HPLC purified, >95%). Protein concentration was determined by spectroscopic method in the presence of 8 M urea (33).

### LLPS imaged by differential interference contrast (DIC) microscopy

The formation of liquid droplets was imaged on 50 μl of the N protein samples by DIC microscopy (OLYMPUS IX73 Inverted Microscope System with OLYMPUS DP74 Color Camera) as previously described (10,14). The N protein samples were prepared at 10 μM in 25 mM HEPES buffer (pH 7.0) with 70 mM KCl (buffer 1), while NTD and CTD samples were prepared at 100 μM and 200 μM respectively in 10 mM sodium phosphate buffer (pH 7.0) in the presence of 150 mM NaCl (buffer 2) for NMR studies. HCQ at 10 mM was dissolved in buffer 1 for for LLPS or buffer 2 for NMR binding studies with the final pH adjusted to 7.0.

### NMR characterizations of the binding of HCQ to NTD and CTD

NMR experiments were conducted at 25 °C on an 800 MHz Bruker Avance spectrometer equipped with pulse field gradient units and a shielded cryoprobe as described previously (10,14,19,22,34). For NMR HSQC titrations with HCQ, two dimensional ^1^H-^15^N NMR HSQC spectra were collected on the ^15^N-labelled NTD at 100 μM or CTD at 200 μM in the absence and in the presence of HCQ at different ratios. NMR data were processed with NMRPipe (35) and analyzed with NMRView (36).

### Calculation of CSD and data fitting

Sequential assignments were achieved based on the deposited NMR chemical shifts for NTD (2) and CTD (37). To calculate chemical shift difference (CSD), HSQC spectra collected without and with HCQ at different concentrations were superimposed. Subsequently, the shifted HSQC peaks were identified and further assigned to the corresponding NTD and CTD residues. The chemical shift difference (CSD) was calculated by an integrated index with the following formula (10,19,20,22):

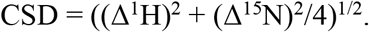

In order to obtain residue-specific dissociation constant (Kd), we fitted the shift traces of the NTD and CTD residues with significant shifts (CSD > average + STD) by using the one binding site model with the following formula as we previously performed (10,19,20,22):

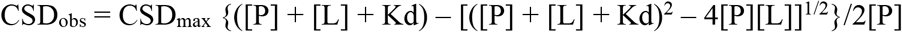

Here, [P] and [L] are molar concentrations of NTD/CTD and ligands (HCQ) respectively.

### Molecular docking

The structures of the HCQ-NTD and HCQ-CTD complex were constructed by use of the well-established HADDOCK software (10, 20,22,23) in combination with crystallography and NMR system (CNS) (38), which makes use of CSD data to derive the docking that allows various degrees of flexibility. Briefly, HADDOCK docking was performed in three stages: (1) randomization and rigid body docking; (2) semi-flexible simulated annealing; and (3) flexible explicit solvent refinement.

The NMR structure (2) of NTD (PDB ID of 6YI3) and crystal structure (3) of CTD (PDB ID of 6YUN) were used for docking to HCQ. The HCQ-NTD and HCQ-CTD structures with the lowest energy score were selected for the detailed analysis and display by Pymol (The PyMOL Molecular Graphics System).

## Acknowledgement

This study is supported by Ministry of Education of Singapore (MOE) Tier 1 Grants R-154-000-B92-114 to Jianxing Song.

## Author contributions

Conceived the research: J.S. Performed research and analyzed data: M.D and J.S; Acquired funding: J.S; Wrote manuscript: J.S.

## Competing interests

The authors declare no competing interests.

## Supplementary Materials

**Table S1.**
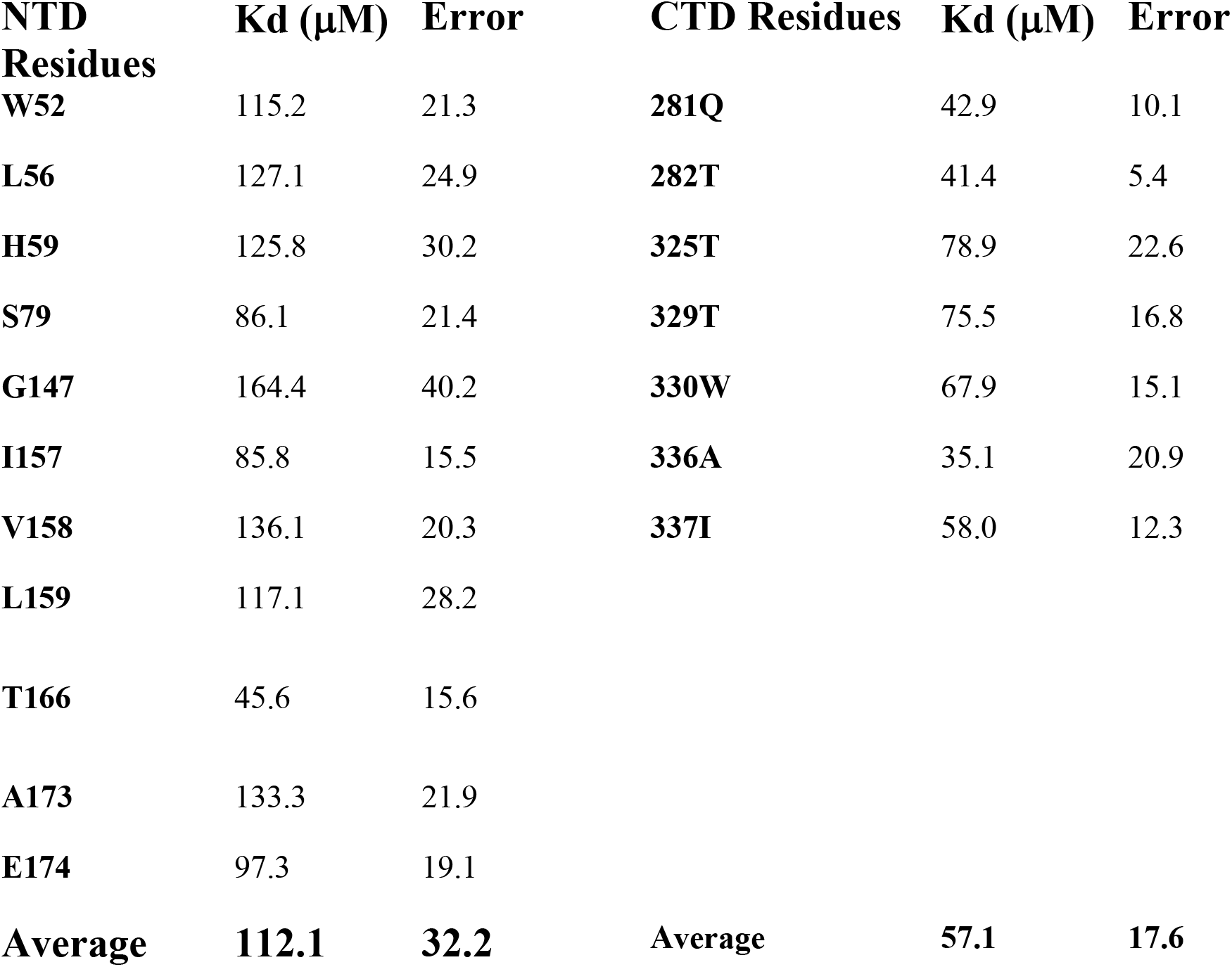
Residue-specific dissociation constants (Kd) of NTD and CTD residues.

**Fig. S1.**
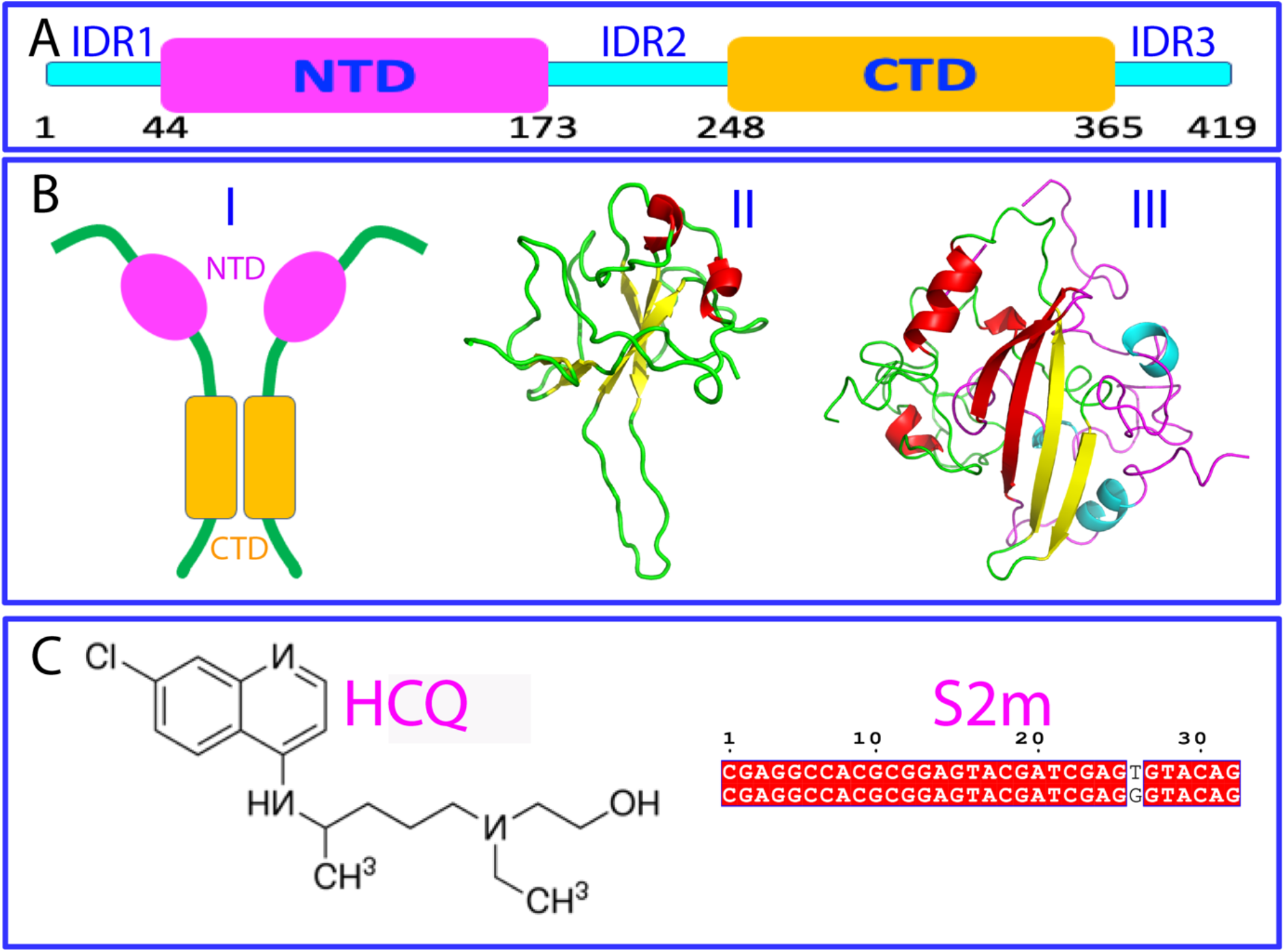
Structures of SARS-CoV-2 N protein, HCQ and S2m. (A) Domain organization of SARS-CoV-2 N protein. (B) Schematic representation of the dimeric N protein (I), three-dimensional structures of NTD (II) and dimeric CTD (III). (C) Chemical structures of hydroxychloroquine (HCQ) and sequences of 32-mer ssDNA S2m of SARS-Cov-1 (upper) and SARS-CoV-2 (lower).

**Fig. S2.**
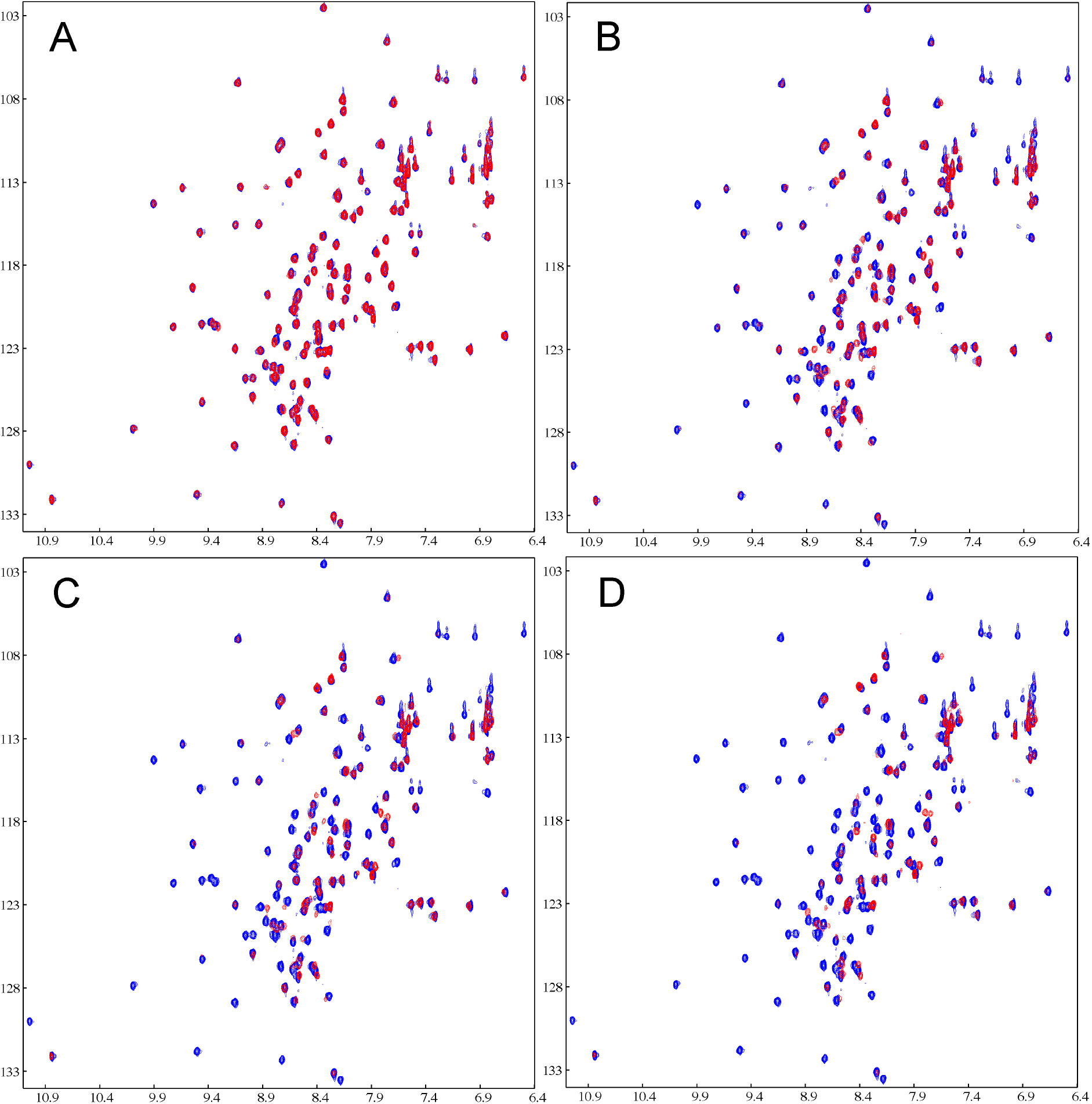
S2m binds NTD of SARS-CoV-2 N protein. Superimposition of HSQC spectra of NTD at 100 μM in the free state (blue) and in the presence of S2m (red) at 1:0.1 (A), 1:0.5 (B), 1:1.0 (C) and 1:2.5 (D) (NTD:S2m).

**Fig. S3.**
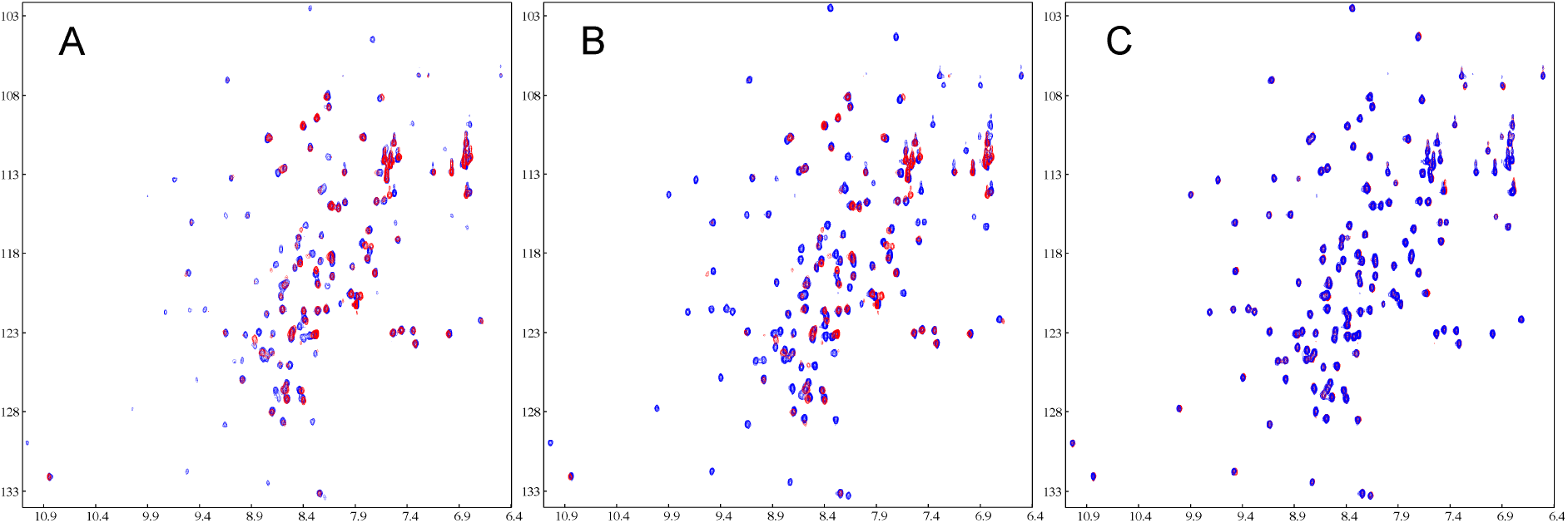
HCQ displaces S2m from binding with NTD of SARS-CoV-2 N protein. Superimposition of HSQC spectra of NTD at 100 μM in the presence of S2m at 1:2.5 (NTD:S2m) (red) and with additional addition of HCQ (blue) at 3.75 (A) and 1:15 (B). (C) Superimposition of HSQC spectra of NTD at 100 μM in the presence of only HCQ at 1:15 (blue) and in the presence of both S2m at 1:2.5 and HCQ at 1:15 (red).

**Fig. S4.**
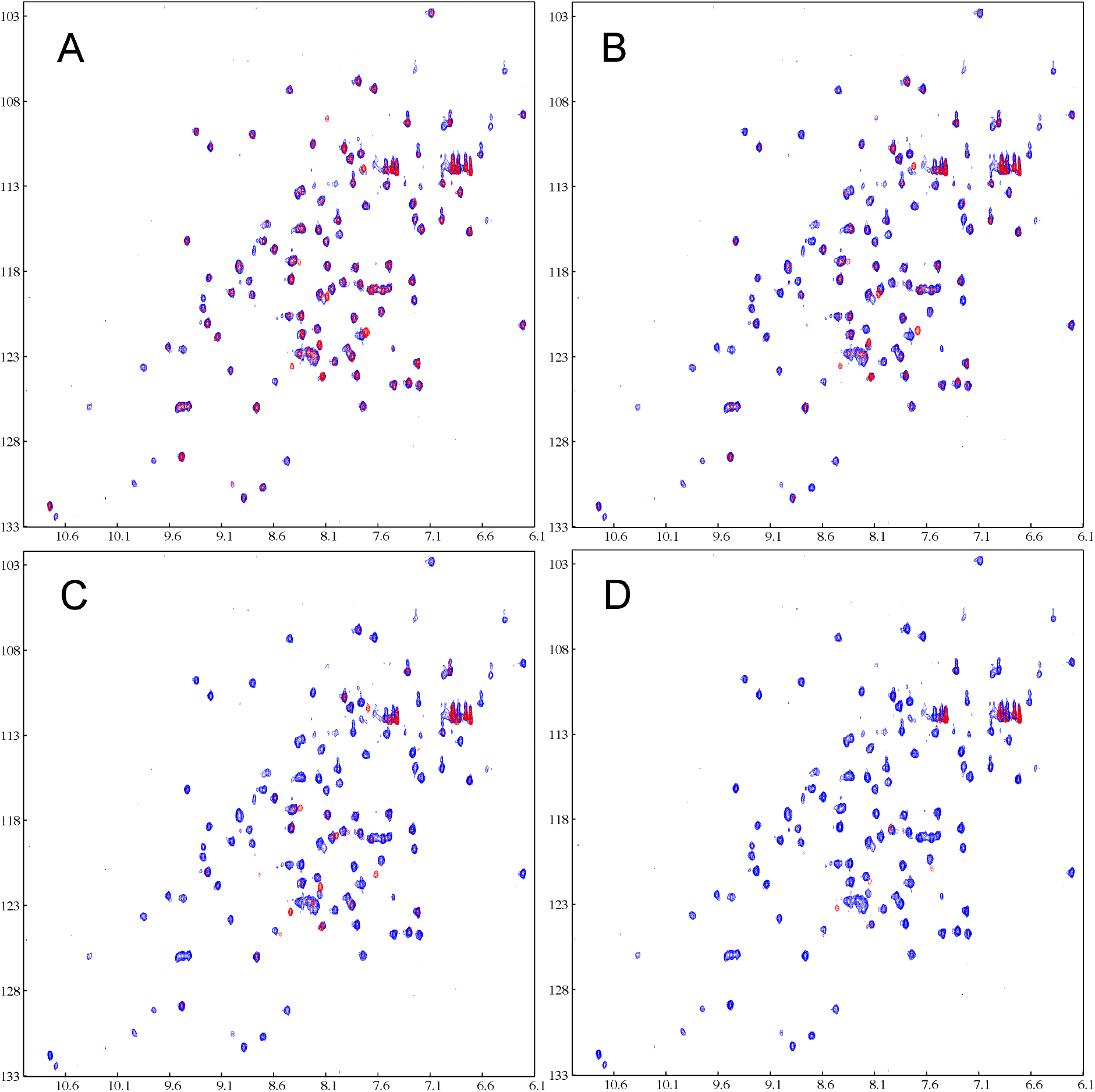
S2m binds CTD of SARS-CoV-2 N protein. Superimposition of HSQC spectra of CTD at 200 μM in the free state (blue) and in the presence of S2m (red) at 0.05 (A), 0.1 (B), 0.5 (C) and 1.0 (D).

**Fig. S5.**
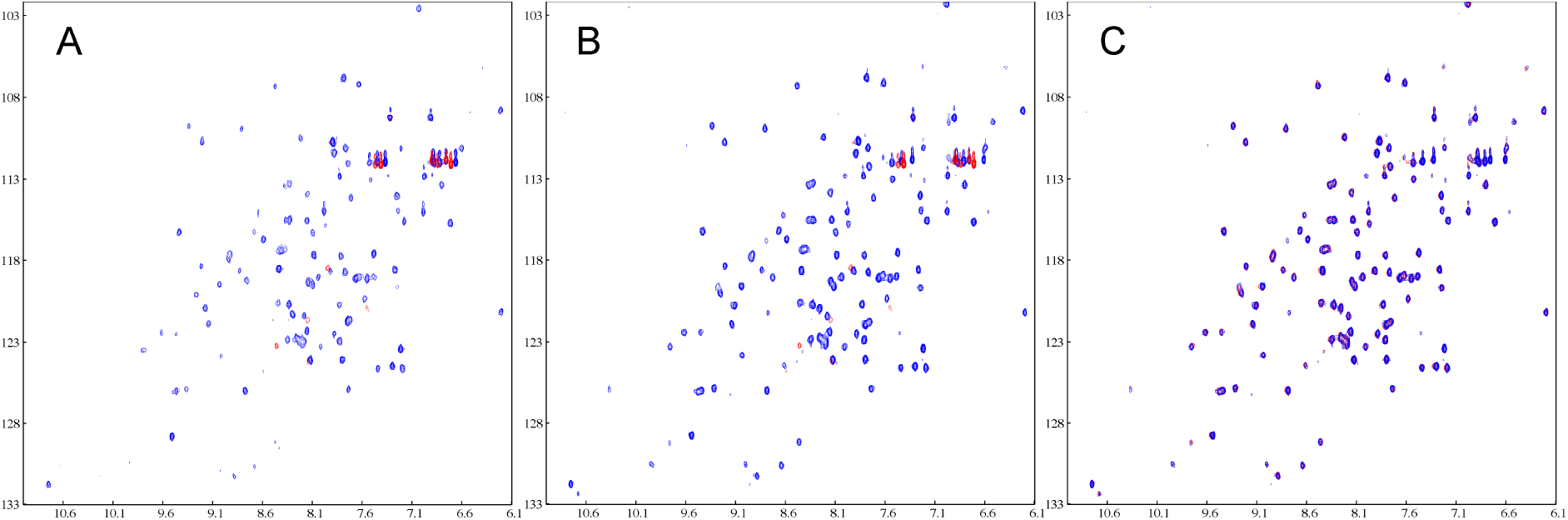
HCQ displaces S2m from binding with CTD of SARS-CoV-2 N protein. Superimposition of HSQC spectra of CTD at 200 μM in the presence of S2m at 1:1 (CTD:S2m) (red) and with additional addition of HCQ (blue) at 1:88 (A) and 1:7.5 (B). (C) Superimposition of HSQC spectra of CTD at 200 μM in the presence of only HCQ at 1:7.5 (blue) and in the presence of both S2m at 1:1 and HCQ at 1:7.5 (red).

**Fig. S6.**
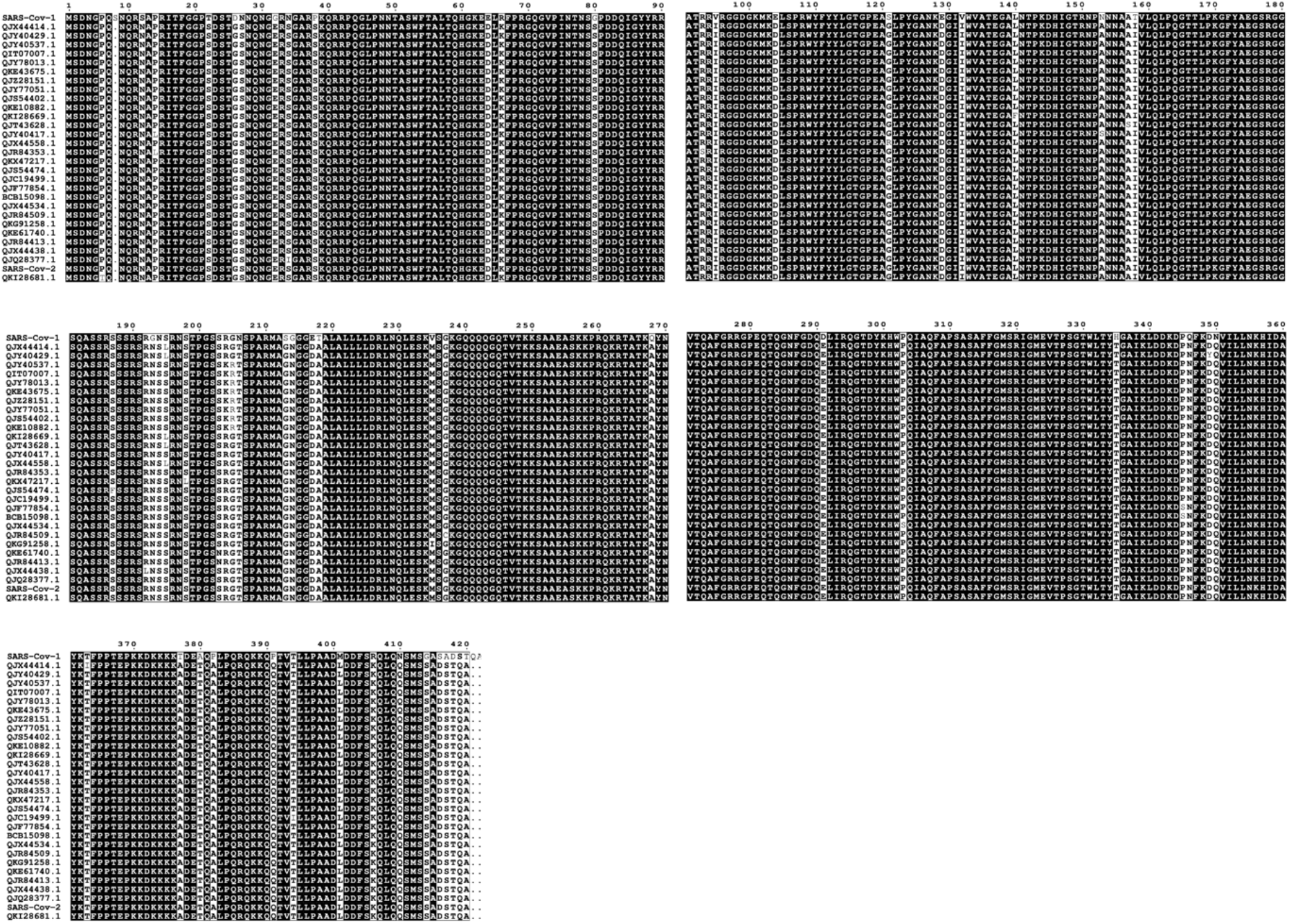
Sequence alignment of nucleocapsid (N) proteins of SARS-CoV-1 as well as SARS-CoV-2 and its major variants.

**Fig. S7.**
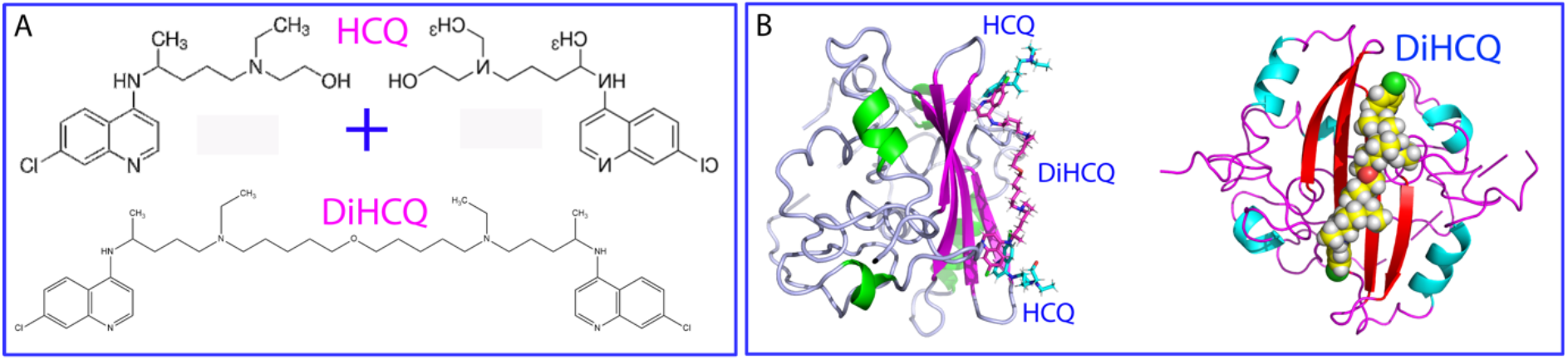
A proposed strategy to design better anti-SARS-CoV-2 molecule from HCQ with the higher affinity and specificity. (A) Schematic representation of linking two HCQ molecules to form DiHCQ. (B) Docking structures of the dimeric CTD in complex with HCQ or with DiHCQ with HCQ/DiHCQ displayed in stick or sphere respectively.

## Notes

### Competing Interest Statement

The authors have declared no competing interest.

### Summary of Updates

1) All detailed experimental results are included. 2) All supplementary materials are included. 3) The manuscript has been finalised into the submitted format.

